# Haploinsufficiency of Parkinsonism Gene *SYNJ1* Contributes to Dopamine neuron Vulnerability in Aged Mice

**DOI:** 10.1101/233585

**Authors:** Ping-Yue Pan, Patricia Sheehan, Qian Wang, Yuanxi Zhang, Jing Wang, Farida El Gaamouch, Li Zhu, Dongming Cai, Zhenyu Yue

**Affiliations:** Neurology Department; The Friedman Brain Institute, Icahn School of Medicine at Mount Sinai, New York, NY 10029 USA; James J Peters VA Medical Center, Research & Development, Bronx, NY 10468 USA

## Abstract

Parkinson’s disease (PD) is an age-dependent neurodegenerative disorder characterized by the loss of substantia nigra dopaminergic (DAergic) neurons in ventral midbrain (MB). Identification of interactions between aging and the known risk variants is crucial to understanding the etiology of PD. Recessive mutations in *SYNJ1* have recently been linked to familial early-onset atypical Parkinsonism. We now show an age-dependent decline of *SYNJ1* expression in the striatum as well as in striatal DAergic terminals of aged mice. Heterozygous deletion of *SYNJ1* in mice causes selective elevation of PIP_2_ in the MB, and manipulation of PIP_2_ levels also impairs synaptic vesicle recycling preferentially in MB neurons. *SYNJ1*^+/−^ mice display progressive PD-like behavioral alterations and DAergic terminal degeneration. Furthermore, we found down-regulation of human *SYNJ1* transcripts in a subset of sporadic PD brains, corroborating the role of an age-dependent decrease in *SYNJ1* in predisposing DAergic neuron vulnerability and PD pathogenesis.

## Introduction

Parkinson’s disease (PD) is one of the most debilitating neurodegenerative disorders affecting millions of people worldwide. Most cases are diagnosed later in life, and the incidence of PD grows exponentially after the age of sixty (Driver et al., 2009). Due to the sporadic nature of most cases, it has been difficult to determine the underlying pathogenic mechanism. Emergent evidence suggested convergent pathogenic pathways, including dysfunctional synaptic membrane trafficking, during disease progression (Trinh and Farrer, 2013; Schirinzi et al., 2017). Human genetic studies and genome-wide association studies (GWAS) have also revealed an overlapping pool of genes, such as LRRK2 and SNCA, that contributes to both familial and sporadic PD (Singleton et al., 2013; Spataro et al., 2015; Hernandez et al., 2016).

*SYNJ1/PARK20* is one of the most recent additions to the *PARK* gene family, as three recessive point mutations – R258Q, A459P, and R839C, in both the SAC1 domain and the 5-phosphatase domain of the *SYNJ1* gene, have been identified in families with juvenile atypical Parkinsonism with epilepsy (Krebs, et al., 2013; Quadri, et al., 2013; Olgiati, et al., 2014; Kirola, et al., 2016; Taghavi, et al., 2017). Synaptojanin1 (synj1, encoded by *SYNJ1)* is an important phosphoinositide phosphatase, which has two isoforms – the 170-kDa isoform a (NP_003886.3) and the 145-kDa isoform b (NP_982271.2). The brain-specific isoform b is enriched in nerve terminals where it regulates synaptic vesicle (SV) recycling and synaptic protein targeting (McPherson et al., 1996; Micheva et al., 1997; Mani et al., 2007; Dong et al., 2015) by hydrolyzing membrane phosphoinositides (e.g. PI(4,5)P_2_, PI(3,4)P_2_, PI3P and PI4P) via its two phosphatase domains and by binding to endophilin at the proline-rich domain. Knock-in mice carrying the homozygous R258Q disease allele, which abolishes the SAC1 activity (Krebs et al., 2013) recapitulated parkinsonian symptoms and exhibited defective clathrin uncoating and dystrophic changes in the nigrostriatal terminals (Cao et al., 2017b). Interestingly, missense mutations in auxilin (encoded by *DNAJC6/PARK19*), another protein involved in clathrin uncoating, were also identified in early-onset familial forms of PD (Edvardson et al., 2012; Koroglu et al., 2013). The pathogenic mechanism connecting dysfunctional synaptic membrane trafficking and PD pathogenesis, however, remains unclear, although a recent study in *Drosophila* suggests a potential link to autophagic deregulation (Vanhauwaert et al., 2017). Nonetheless, disease mutations in familial PD represent rare occurrences in the affected population. A clinical study has now found variations in the non-coding regions of *DNAJC6* in sporadic PD cases (Olgiati et al., 2016). More strikingly, in a recent meta-analysis for the sporadic PD GWAS, the gene encoding endophilinA, *SH3GL1*, was revealed as one of 17 new risk variants (Chang, et al., 2017). These results have brought increasing interest in understanding if synj1, a close interacting partner of endophilinA, contributes to risks in sporadic PD via common pathways at the nerve terminal.

For sporadic PD, aging is considered the greater contributor compared with genetic risks (Driver et al., 2009; Collier et al., 2011). Major pathogenic pathways implicated in PD such as mitochondrial dysfunction, membrane trafficking and autophagy-lysosomal impairment, increased oxidative stress and neuroinflammation also deteriorate during normal aging. More importantly, the number of substantia nigra *par compacta* (SNpc) DAergic neurons in the ventral midbrain (MB), whose degeneration is known as the hallmark of PD, is also markedly reduced in healthy aged subjects (Stark and Pakkenberg, 2004; Mortera and HerculanoHouzel, 2012). How aging interacts with genetic variants and triggers the clinically significant pathological course of PD remains to be understood.

To understand whether loss of function of *SYNJ1* leads to DAergic vulnerability in aging-related PD pathogenesis, we examined the impact of aging on *SYNJ1* expression and performed a detailed investigation in the previously established *SYNJ1*^+/−^ mice (Voronov, et al., 2008). We demonstrate an age dependent decline of *SYNJ1* expression in the striatum, particularly in the DAergic terminals; and age-dependent reduction in *SYNJ1* predisposes *SYNJ1*^+/−^ mice to PD-like motor dysfunctions and striatal DAergic degeneration. Our study demonstrates for the first time haploinsufficiency of a Parkinsonism gene contributes to dopamine neuron vulnerability in aged mice.

## Results

### *SYNJ1* transcript levels are reduced in the striatum of sporadic PD brains and healthy aged brains

To understand the role of *SYNJ1* in sporadic PD, we visited data sets in the public domain (GSE28894, GSE20168, GSE8397) and found down-regulation of *SYNJ1* transcripts in multiple brain regions including the prefrontal cortex, the striatum and the substantia nigra (SN) in subsets of sporadic PD brains (Figure 1A). The reduction of *SYNJ1* in these brain regions, except the lateral SN, is unlikely due simply to neuronal or synaptic loss, as the synaptic marker, *SYP* (encoding synaptophysin) was largely unaltered in the same subjects (Figure 1B). This result encouraged us to explore the potential link between *SYNJ1* loss-of-function and age-dependent sporadic PD. To investigate the impact of aging on *SYNJ1* expression, we first examined postmortem human data from public databases by extracting the raw measurements of *SYNJ1* transcripts in developing human brains (data from Allen Brain Atlas, lacking SN data) and in aged human brains (data from GTEx) (see **Materials and Methods / human data analysis**). To better elucidate age-dependent changes in transcript levels, we binned the data by age groups and normalized both data sets (developing and aged) to the mean transcript level in the cortex of the overlapping age group 21-30 (Figure. 1C left). An accelerated decline of *SYNJ1* transcripts was observed in the striatum from 11-20 years (bin 3) to 41-60 years (bin 6), while cortical *SYNJ1* remained largely steady. The decline of *SYNJ1* in the striatum is not due to neuronal loss or synapse loss as β-actin (Figure 1C middle) and synapsin1 (Figure 1C right) levels do not exhibit similar trends in the striatum. However, the decrease of *SYN1* (synapsin 1) expression became apparent at 61-70 years in the cortex, and was accompanied by a comparable reduction in the *SYNJ1* level, suggesting synapse loss. The age-dependent down-regulation of *SYNJ1* in the SN is less clear due to lack of available data, although the average values in most binned groups appear similar to those in the striatum.

**Figure 1.**
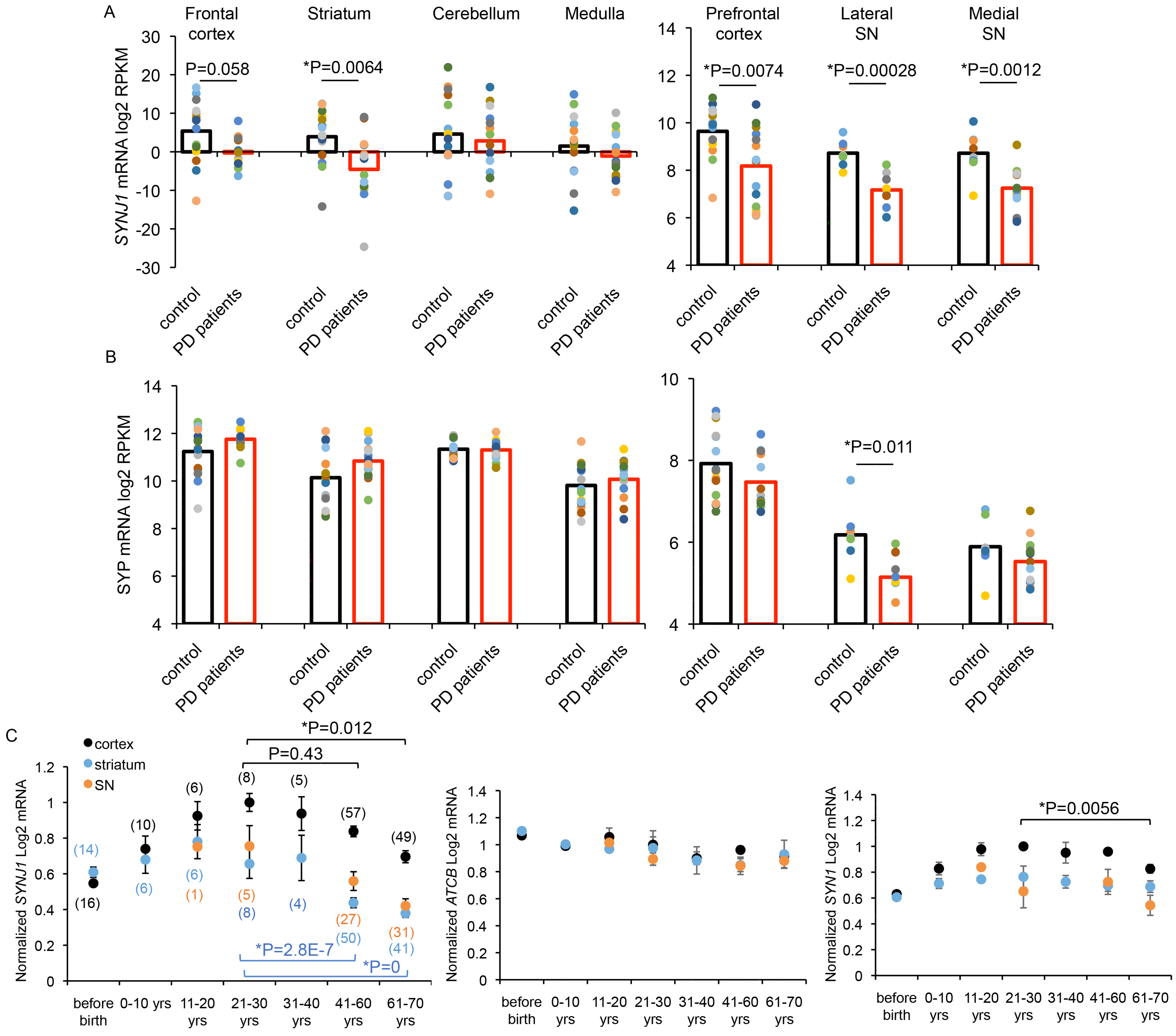
*SYNJ1* transcripts are reduced in subsets of sporadic PD brains and the striatum of aged humans. **A-B)** *SYNJ1* (**A**) and *SYP* (synaptophysin) (**B**) mRNA levels from postmortem sporadic PD and control human samples were analyzed based on datasets GSE28894, GSE20168, GSE8397 (see Materials and Methods / Human Data Analysis). Each data point represents one patient. P values are from two-sample Student’s *t* test. **C**) *SYNJ1* (synj1, top), *ACTB* (β-actin, middle) and *SYN1* (synapsin I, bottom) transcript levels in different brain regions were normalized to the average of the “21-30 years” bin and plotted against all binned age groups. Data sets obtained from Allen Brain Atlas developing human brain and GTEx aged brain bank. Numbers in brackets indicate the number of subjects in each binned age group. To test the null hypothesis that *SYNJ1* expression is not down-regulated by aging, we performed one-way ANOVA followed by Tukey’s *post-hoc* tests for each brain region starting from bin 3, 11-20 years old, to bin 7, 60-70 years old. One-way ANOVA for *SYNJ1:* cortex, *P = 4.52E-4; striatum, *P = 0; MB, *P = 0.014. One-way ANOVA for *SYN1:* cortex, *P = 0.0059; striatum, P = 0.95; MB, P = 0.57. P values are from Tukey’s *post-hoc* analysis for brain regions display significant age-dependent changes.

To verify the above findings in an animal model and to determine if the decline of *SYNJ1* is reflected beyond the mRNA level to the protein level, we performed immunohistochemical analyses of C57/BL6 mouse brains of both sexes at various ages. We first verified the specificity of the synj1 antibody for its synapse-enriched localization and its validity for quantitative measures in both cell cultures and brain slices (Figure 1-Figure supplement 1). We then performed immunohistochemical staining for tyrosine hydroxylase (TH), synj1, and synapsin1/2, a presynaptic marker, in coronal sections that included both the striatum and the cortex (Figure 1-Figure supplement 2A). The immunofluorescence of synj1 and synapsin1/2 in the striatum was normalized to that in the cortex for each slice (see **Materials and Methods/Data analysis**). While synapsin1/2 level remained at a constant 80% relative to the cortex across all age groups, synj1 expression was significantly reduced in the striatum of 18-month old mice (Figure 1-Figure supplement 2B), reminiscent of the aged human brain.

### Synj1 expression in striatal DAergic terminals is reduced in aged mice

The striatum is an essential part of the basal ganglia, where DAergic terminals from the ventral MB and glutamatergic (GLUTergic) terminals from the cortex converge at striatal medium spiny neurons to regulate motor output (Figure 2A). To determine if DAergic terminals in the striatum also exhibit an age-dependent change in synj1 expression, we performed an in-depth analysis of the immunofluorescence in the coronal sections from 3-month and 18-month old mice. By colabeling with TH, the rate-limiting enzyme for DA synthesis, and synapsin1/2, a presynaptic marker, we were able to differentiate DAergic terminals (TH and synapsin1/2 positive) and non-DAergic terminals (TH negative, synapsin1/2 positive). We found that in DAergic terminals, synj1 expression was 10% lower than in non-DAergic terminals at 3-month old (Figure 2B-K); however, this difference grew to nearly 50% in 18-month old animals. Consistently, the cumulative probability for synj1 immunofluorescence sampled from over 1000 nerve terminals revealed a greater difference between TH^+^ and TH^-^ terminals in an 18-month old mouse compared to a 3-month old mouse (Figure 2 E, K).

**Figure 2.**
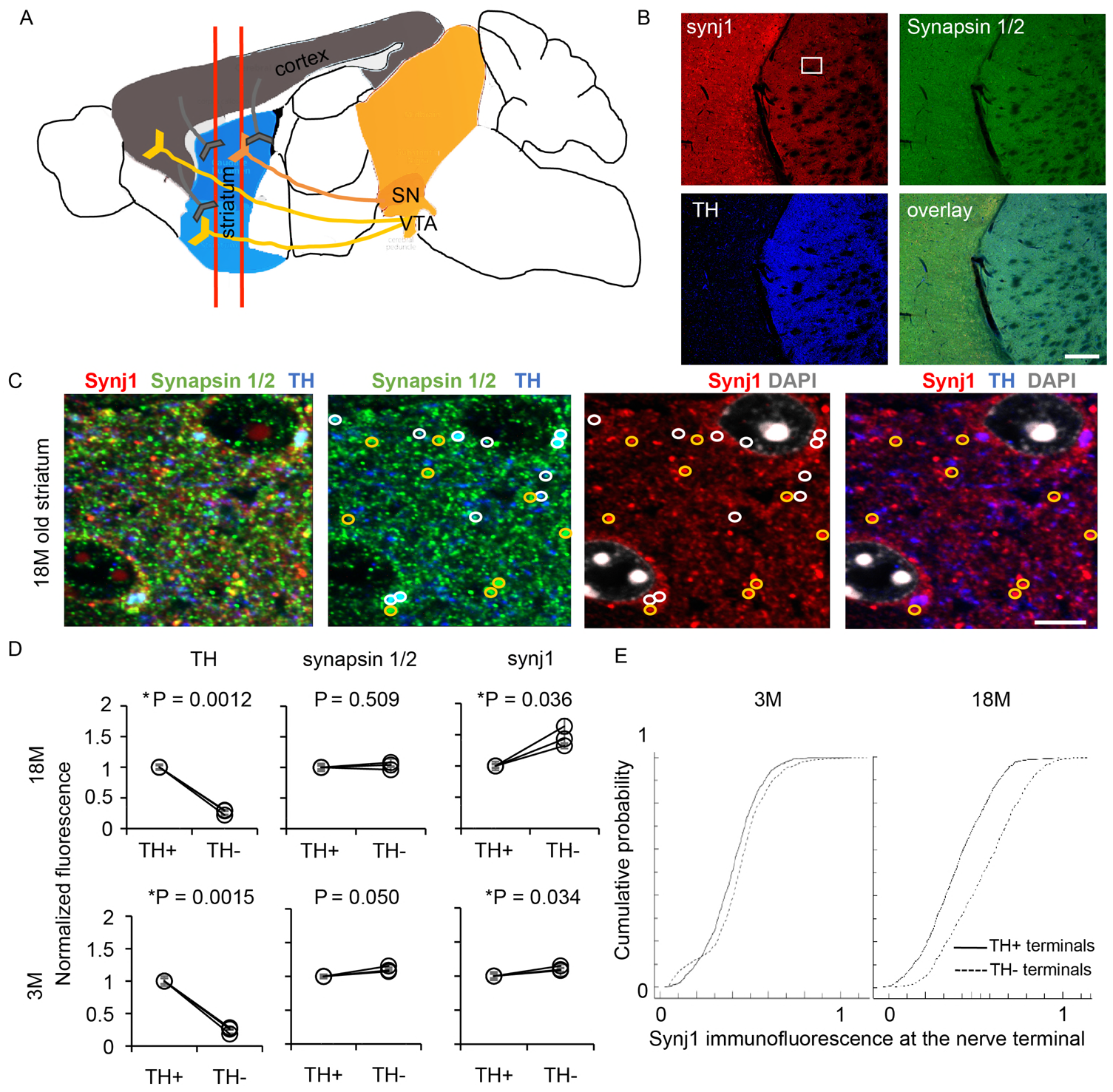
Synj1 immunoreactivity is reduced in striatal dopaminergic terminals in aged mice. **A**) Diagram illustrating positions where 40 nm cryoslices were taken from the mouse brain **B-C**) Low-magnification (**B**) and High-magnification (**C**) confocal images of a representative 18-month old striatal slice immunolabeled with anti-TH (blue), anti-synj1 (red) and anti-synapsin1/2 (green). White and yellow circles in **C** represent TH-positive (DAergic) and TH-negative nerve terminals, respectively. Scale bar in **B**, 200 μm, scale bar in; **C**, 5μm. **D**) Comparison of averaged TH, synapsin1/2 and synj1 immunoreactivity in circular regions of interests (as in **C**) for 3-month and 18-month old mice. Each data point represents the average of the mean fluorescence measured from 10-20 images of 2 slices for each mouse and the error bar is the standard error represents each of the 3 mice. P values are calculated by one-sample Student’s *t* tests (see Materials and Methods / Data analysis). E) Cumulative probability distribution for synj1 immunoreactivity at TH^+^ and TH^−^ terminals in a mouse at 18-month old (TH^+^ terminals: N = 694; TH^−^ terminals: N = 985, D = 0.25, *P < 0.001, two-sample Kolmogorov-Smirnov test) and another at 3-month old (TH+ terminals: N = 1021, TH^−^ terminals: N = 1011, D = 0.15, *P < 0.001, two-sample Kolmogorov-Smirnov test).

### MB displays specific vulnerability to PIP_2_ metabolism in *SYNJ1*^+/−^ mice

The primary function of synj1 is to regulate phosphoinositide metabolism and support the normal functions of membrane trafficking. To understand the impact of synj1 expression on membrane phosphoinositide levels, we measured the content of PIP_2_, PIP and PI in the cortex, the striatum and the MB of 1-year old *SYNJ1*^+/−^ mice and littermate wildtype (WT) mice using high-performance liquid chromatography (HPLC). We first noted that the PIP_2_ level in the MB is nearly 2-fold higher, whereas the PI and the PIP levels were significantly lower, than those in the cortex (n=23) (Figure 3A). The PIP_2_ level in the MB was further elevated by 15% in 1-year old *SYNJ1*^+/−^ mice than that of WT (p=0.007) (Figure 3B). While there is a trend of increased PIP_2_ in the striatum (p=0.07), no obvious change in the cortex of *SYNJ1*^+/−^ mice was observed (Figure 3B). The amount of PIP_2_ increase in the heterozygous MB is commensurate to a previous study, which found PIP_2_ levels to be elevated by approximately 12% in the whole brain samples of *SYNJ1*^+/−^ mice (Voronov, et al., 2008).

**Figure 3.**
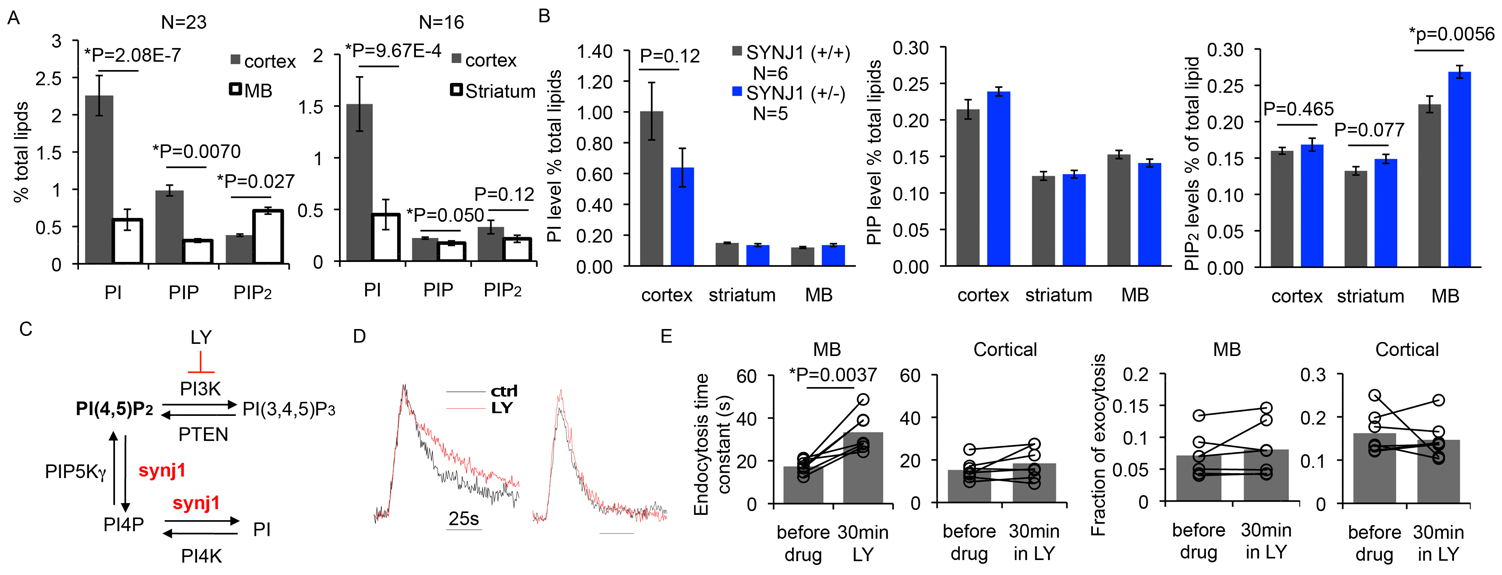
MB-specific sensitivity to PIP_2_ deregulation. **A**) Comparison of phosphoinositide contents measured by HPLC in different brain regions (cortex, MB and striatum) of male WT mice at 12 months old. **B**) Summary of PI, PIP and PIP_2_ levels from 12 months old *SYNJ1*^+/−^ mice and *SYNJ1*^+/+^ mice of both sexes. P values are calculated by two-sample Student’s *t* tests. **C**) Illustration for the role of synj1 in regulating phosphoinositide metabolism and the role of LY249002 (LY) in inhibiting the conversion of PI(4,5)P_2_ to PI(3,4,5)P_3_ **D**) Representative pHluorin signal for the cultured MB and cortical neurons at 10 Hz, 10 s stimulation before (black) and after LY incubation (red). **E**) Summary of endocytosis and exocytosis kinetics before and after LY treatment in different neurons. P values are calculated by paired Student’s *t* tests.

To determine how PIP_2_ accumulation in the MB could affect neuronal function, we examined the efficiencies of SV recycling in cultured neurons by expressing an optical reporter, pHluorin. PHluorin is a pH-sensitive variant of GFP whose protonation and deprotonation results in a dynamic 20-fold change in fluorescence, which allows for the quantitative measurement of SV exocytosis and endocytosis when conjugated to the lumenal aspect of the vesicular transporter (Sankaranarayanan et al., 2000; Ariel and Ryan, 2010; Pan and Ryan, 2012). VMAT2-pHluorin or vGLUT1-pHluorin was expressed in cultured MB or cortical neurons, respectively, and a 10 Hz, 10 s field stimulation was applied to trigger SV recycling. We previously showed that cultured MB but not cortical neurons from *SYNJ1*^+/−^ mice displayed slowed endocytosis (Pan et al., 2017), suggesting that insufficient conversion of PIP_2_ due to heterozygous deletion of *SYNJ1* affects SV endocytosis preferentially in MB neurons. To verify the selective effect of altered PIP_2_ levels in MB neurons, we treated cultured neurons with LY249002 (LY), a PI3K inhibitor, which blocks the conversion of PIP_2_ to PIP_3_ on the plasma membrane, and found that SV endocytosis in MB neurons was substantially slower after a 10-15 min treatment (Figure 3C-E). In contrast, cortical neurons showed minimal responsiveness to LY incubation up to 30 min. The changes in the amount of exocytosis in the MB neurons are more heterogeneous across different nerve terminals and do not exhibit any difference on average. Our data suggests that SV trafficking in MB neurons is more susceptible to PIP_2_ accumulation than cortical neurons.

### *SYNJ1*^+/−^ mice display PD-like motor function deficits

We next evaluated the motor functions associated with the reduced expression of *SYNJ1.* Mice with complete deletion of *SYNJ1* are not viable and die shortly after birth. *SYNJ1*^+/−^ mice, however, appear normal with regard to body size and life span. Unlike the *R258Q* disease mutation homozygous knock-in mice (*RQ KI*), some of which display severe movement problems and tonic-clonic seizures (Cao et al., 2017b), *SYNJ1*^+/−^ mice do not have seizures or apparent gait difficulties in their lifetime. When *SYNJ1*^+/−^ and *SYNJ1*^+/+^ littermates were evaluated for their general locomotor functions in the open-field assay, 7-month old *SYNJ1*^+/−^ mice exhibited hyperactivity (Figure 4A) compared to their littermates. Both *SYNJ1*^+/+^ and *SYNJ1*^+/−^ mice displayed a significant deterioration in their activity levels at 12-months old and *SYNJ1*^+/−^ mice appeared no different than WT mice at this age (Figure 4B). To further test the integrity of dopamine-regulated motor function, we challenged the mice with quinpirole, a DA D2 receptor (D2R) agonist. Motor inhibition induced by quinpirole was significantly impaired in *SYNJ1*^+/−^ mice at 12-months old (Figure 4C), as was their motor coordination when tested on the accelerated Rota-rod (Figure 4D). Such decline in motor functions following hyperactivity in *SYNJ1*^+/−^ mice is reminiscent of the findings in many other PD models (Chesselet and Richter, 2011, review), including the *LRRK2 G2019S KI* mice (Volta, et al., 2017).

**Figure 4.**
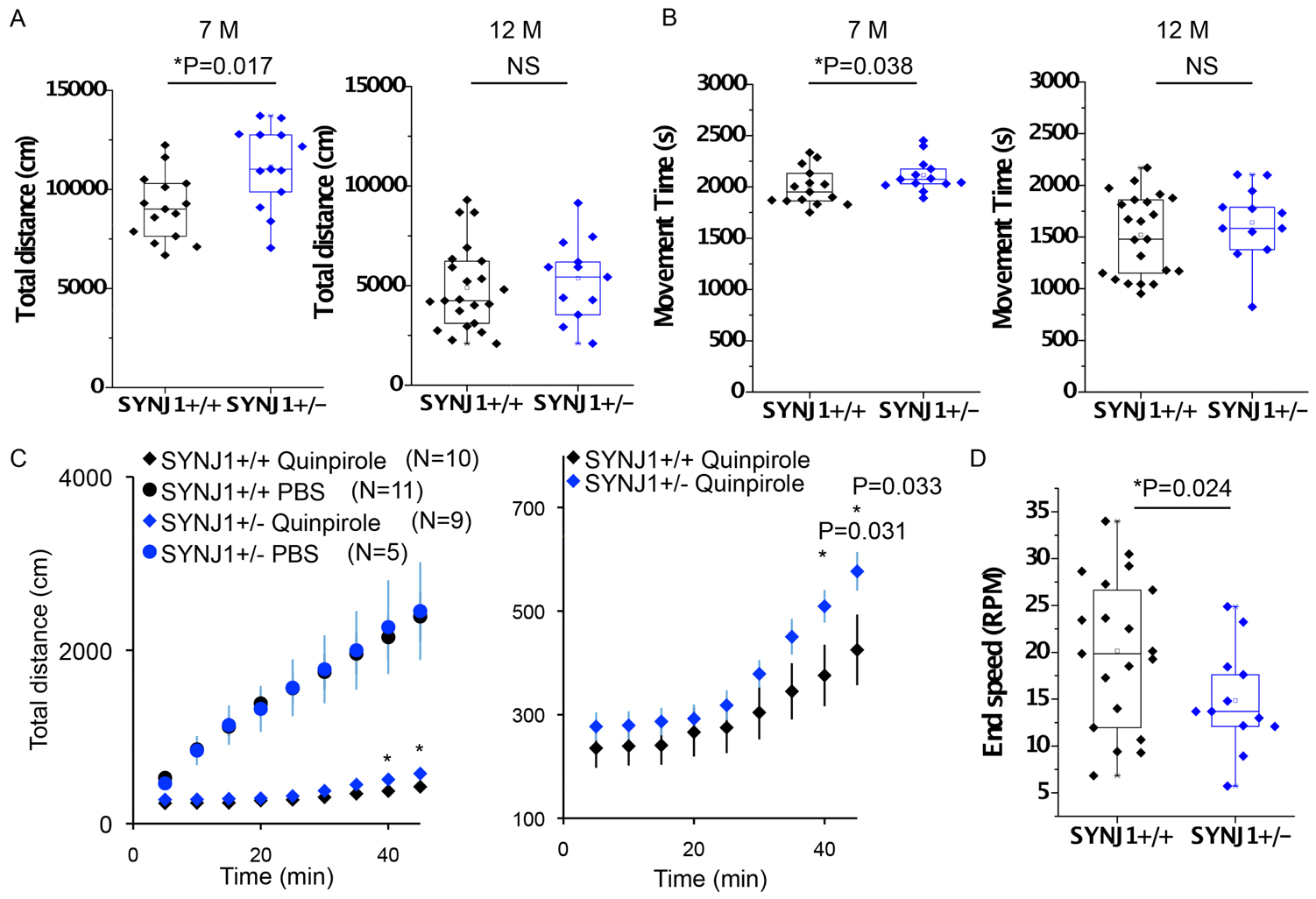
*SYNJ1*^+/−^ mice exhibit hyperactivity in early adulthood and motor function alterations. **A-B**) Total distance (**A**) and movement time (**B**) measured in the open-field assay for male *SYNJ1*^+/+^ and *SYNJ1*^+/−^ mice at 7-month and 12-month of age, respectively. A cohort of *SYNJ1*^+/+^ (N=15) and *SYNJ1*^+/−^ (N=13) littermates were examined at 7 months old; and a cohort of *SYNJ1*^+/+^ (N=22) and *SYNJ1*^+/−^ (N=13) littermates were examined at 12 months old. Due to an imbalance in the number of subjects, two-way repeated measures ANOVA with *post-hoc* tests do not apply. P values are calculated by two-sample Student’s *t* tests. **C**) Total distance in the open field assay for 12 months old mice treated (IP injection) with PBS or quinpirole (0.5 mg/kg) – a D2 receptor agonist. Locomotor activity was averaged within each 5-minute bin of recording and 12 data points were obtained for each animal for a total of 60 minutes of recording. Error bars represent standard errors for the number of animals tested at each 5-minute bin. P values are from two-sample Student’s *t* tests. **D**) Accelerated Rota-rod assay summarizing the end speed before mice fell off the rotating bar. P values are from two-sample Student’s *t* tests.

### Aged *SYNJ1*^+/−^ mice exhibit loss of DAergic terminals

To further understand the pathological consequence of reduced *SYNJ1* expression, we performed the following analyses: 1) the integrity of DAergic neurons and their nerve terminals 2) striatal DA content and metabolism. Stereological analysis of DAergic neurons in the ventral MB revealed no difference in the number of DAergic cell bodies in aged (18-months old) *SYNJ1*^+/−^ mice (Figure 5-Figure supplement 1). In *SYNJ1*^+/+^ mice, the number of DAergic terminals in striatal slices was reduced by nearly 50% as mice aged from 3-months to 18-months. While the number of DAergic terminals was unaltered in *SYNJ1*^+/−^ mice compared to *SYNJ1^+/+^* at 3-months old, aging led to a significantly exacerbated reduction of an additional 50% in 18-month old *SYNJ1*^+/−^ mice (Figure 5A-B). Consistently, striatal DA content and DA metabolites, measured by HPLC, were also reduced in the *SYNJ1*^+/−^ mice (Figure 5C), insinuating that *SYNJ1* haploinsufficiency leads to the decline of DA release, the loss of DAergic terminals, or a combination of both due to *SYNJ1* haploinsufficiency.

**Figure 5.**
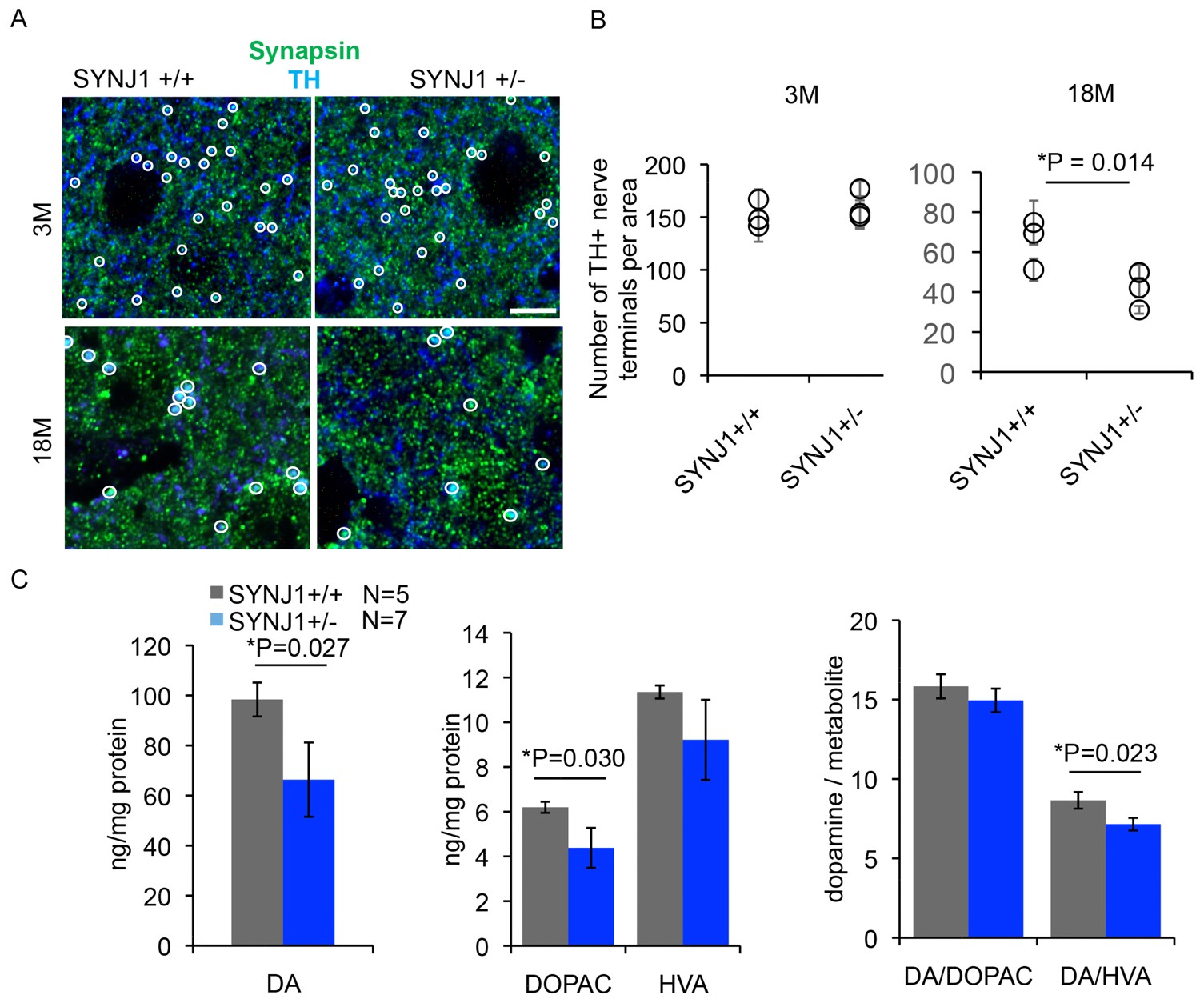
Aged *SYNJ1*^+/−^ mice exhibit loss of dopaminergic terminals and reduction of DA content. **A**) Representative 3-month old and 18-month old striatal slices from a *SYNJ1*^+/+^ mouse and a *SYNJ1*^+/−^ mouse immunolabeled with anti-TH and anti-synapsin1/2. White circles represent DAergic terminals with immunoreactivity for both TH and synapsin1/2. Scale bar, 5 μm. **B**) Summary for the number of DAergic terminals analyzed from 3 male mice. Each data point represents the mean and the standard error for the number of DAergic terminals sampled from 10-20 images for an individual mouse (see Materials and Methods / Data analysis). P value is from two-sample Student’s *t* tests C) DA content and its metabolites levels in the striatum of 12 months old male mice measured by HPLC. P values in both B and C are calculated by two-sample Student’s *t* tests.

## Discussion

Emerging evidence has demonstrated a link between recessive point mutations in *SYNJ1/PARK20* and familial early-onset atypical Parkinsonism. We now show that a reduction of *SYNJ1* expression, which results in aberrant accumulation of PIP_2_ in specific brain regions or neuron populations, may contribute to the risks for age-related sporadic PD. *SYNJ1* heterozygous mice are unable to maintain the proper function of the midbrain DAergic system, driving PD related pathological processes in aged individuals. Our data, therefore, implicates a role of *SYNJ1* loss-of-function in the pathogenesis of age-dependent sporadic PD.

We show that *SYNJ1* expression is down regulated in the striatum during aging in both humans and mice. This is important evidence supporting *SYNJ1* functional insufficiency in PD risk, as most pathogenic changes, such as neuroinflammation and mitochondrial malfunction, are also found to present in normal aging. Whether age-related reduction of *SYNJ1* interacts with other sporadic PD variants has yet to be determined. We previously reported a potential genetic interaction between *LRRK2* disease mutation *G2019S* and *SYNJ1* haploinsufficiency in mice and found LRRK2 mediates phosphorylation of synj1, which leads to an impairment in SV trafficking in MB neurons (Pan et al., 2017). In addition, although several large-scale genome-wide studies failed to reveal *SYNJ1* at the level of significance examined, the gene encoding its closest interacting partner, SH3GL1/endophilinA, is now suggested to be a significant PD-risk gene. Recent studies have also shown that LRRK2 phosphorylates endophilinA and synj1 (Matta et al., 2012; Arranz et al., 2015; Islam et al., 2016; Pan et al., 2017), further suggesting the potential involvement of the LRRK2-endophilinA-synj1 complex in regulating synaptic membrane trafficking, a process which may go awry in the pathogenesis of PD.

While nearly all reported mouse models with complete knockout of recessive PD genes fail to show any PD related phenotypes, our study demonstrates for the first time that haploinsufficiency of a Parkinsonism gene is sufficient to cause DA neuron vulnerability in aged mice. *SYNJ1*^+/−^ mice display PD-like behavioral and pathological changes. *SYNJ1*^+/−^ mice show hyperactivity, which was found in younger mice followed by reduced motor coordination and D2R sensitivity at mid-age. Although it remains to be understood whether and how increased motor activity at younger ages result in a faster decline in motor functions during aging, this phenotype is often found in other PD mouse models, such as the WT and A53T α-synuclein transgenic mice (Unger et al., 2006; Lam et al., 2011; Chesselet et al., 2012) as well as the *LRRK2 G2019S KI* mice (Volta, et al., 2017). Furthermore, aged *SYNJ1*^+/−^ mice display advanced DAergic denervation in the striatum accompanied by reduced DA and DA metabolite content. Interestingly, the combined functional decline and pathological DAergic degeneration are mostly found in mouse models carrying variants with early disease onset and high clinical penetrance (Tsika et al., 2014; Sumi-Akamaru et al., 2015; Cao et al., 2017b). Therefore, our results indicate that *SYNJ1*^+/−^ mice may be used as a potential PD model for dissecting pathogenic pathways at the early stages of PD.

The key question is how reduced levels of *SYNJ1* contribute to vulnerability selectively within the DAergic system. We found that DAergic terminals express less synj1 compared to neighboring non-DA terminals in the striatum. We also showed that cultured ventral MB neurons, including nigral DAergic neurons that project to innervate the striatum, exhibit higher vulnerability to SV recycling compared to cortical neurons in response to *SYNJ1* deficiency (Pan et al., 2017) and significant PIP_2_ elevation. In fact, the elevated levels of PIP_2_ was robust in MB of *SYNJ1*^+/−^ mouse, but not in the cortex, suggesting a lack of compensatory mechanism to maintain the homeostasis of PIP_2_ levels in MB. These impairments could expedite local mechanisms for synapse elimination or axon degeneration (Stevens et al., 2007). Our study did not differentiate DAergic from GABAergic neurons or VTA from SNpc neurons in the ventral MB with respect to their age-dependent change in synj1 levels or their sensitivity to PIP_2_ elevation. Considering the lack of difference in SV endocytosis between TH^+^ and TH^−^ neurons of the ventral MB at baseline (Pan et al., 2017), it is likely that disease protective mechanisms for VTA and GABAergic neurons arise from signaling pathways other than synj1 and lipid alterations. For example, different calcium burdens or neuroinflammatory responses (Pan et al., 2012; Sulzer et al., 2017; Surmeier et al., 2017) may account for their outcome. It was recently reported that loss of synj1 SAC1 activity leads to impaired autophagosome maturation in flies due to abnormal accumulation of PI3P (Vanhauwaert et al., 2017). The PIP levels, however, are not altered in the cortex, the MB or the striatum of the *SYNJ1*^+/−^ mouse in our study. It remains to be tested if changes in PIP_2_ levels alter autophagic signaling via alternative pathways (George et al., 2016) in *SYNJ1*^+/−^ mice that contribute to DAergic neurodegeneration in the striatum.

Interestingly, the SAC1 domain of synj1 that predominantly hydrolyzes phosphatidylinositol monophosphates is considered a weaker enzyme compared to the 5-phosphatase domain, which mainly hydrolyzes PI(4,5)P2 to produce PI4P; yet, the ablation of the SAC1 activity leads to DAergic degeneration in multiple model systems (Cao et al., 2017b; Vanhauwaert et al., 2017). Although the functional outcome of the disease-related mutation, R839C, in the 5-phosphatase domain of synj1 has not been examined; other mutations and variants that were known to substantially impair 5-phosphatase activity were found in patients with early-onset generalized neurological degeneration and epilepsy (Dyment, et al., 2015; Hardies, et al., 2016). The combined clinical and animal data prompts an interesting possibility that reduced expression level or overall function of synj1, rather than the impairment of an individual functional domain is relevant to the pathogenesis of sporadic PD.

Taken together, our findings demonstrate that aging may predispose certain human populations to PD risk via a reduction of *SYNJ1* levels in the striatum. Our study thus not only assists in the identification of novel biomarkers for PD, but also suggests a therapeutic idea for PD by restoring *SYNJ1* levels.

## Materials and methods

### Human data analysis

Sporadic PD brain transcriptome data was downloaded from PMID: 20926834. Among the 17 genome-wide expression datasets, we only examined datasets with a sample size greater than 15 in each group to ensure the statistical power. Three of these datasets, GSE28894, GSE20168, GSE8397, were found to exhibit statistical difference (P < 0.05, two-sample Student’s *t* test) in *SYNJ1* levels in multiple brain regions. Clinical information of the postmortem brain tissue samples can be found in the following articles: PMID 15965975 and 16344956. Normal human brain age-dependent data was downloaded from Allen Brain Atlas / developing human brain (7D4BTI1R5K11_0log2, JOYAOQ1MXT11_0_SYN1_log2, and N2884L1TVT11_3_ACTB_log2, referred to as "Allen brain data” hereafter) which contains data from postconceptional week 8 to 40 years old; as well as from the Genotype-Tissue Expression (GTEx) project, which contains brain region-specific data from 20-70 years old (www.gtexportal.org, referred to as GTEx data hereafter). Raw mRNA data from different brain regions was expressed in the Log2 reads per kilobase per million (RPKM) scale. For Allen brain data, measurements from various parts of the cerebral cortices were averaged and expressed as "cortical” mRNA for each subject. Data for the striatum was collected as a single entry and data for substantia nigra was absent. For each documented age, the number of subjects varied from 1 to 3 in the Allen brain data. For GTEx brain data, a single entry was found for cortex, putamen/basal ganglia (denoted as striatum) and substantia nigra. Data distribution was clustered in the 40-70 years range where typically over 10 subjects were measured for a specific age. We used cortical expression in the 21-30 years age bin for normalization to conjoin the two data sets and reveal age-dependent changes in the full spectrum, but have inevitably lost absolute quantitative information on the log2 scale.

### Animals

Mice were housed in the pathogen-free Center for Comparative Medicine at The Icahn School of Medicine at Mount Sinai. Handling procedures were in accordance with the National Institutes of Health guidelines and approved by the Mount Sinai Institutional Animal Care and Use Committee (IACUC).

### Cryostat, Immunofluorescence and antibodies

Mice were anesthetized with ketamine (100mg/kg) and xylazine (10mg/kg) diluted in saline and perfused transcardially with 4% fresh paraformaldehyde, and post-fixed with 4% paraformaldehyde for over two hours. Dissected brains were cryoprotected in 30% sucrose prior to flash-freezing in the OCT-compound media (SAKURA). Coronal sections were sliced at 40 μm thickness on a Leica CM 3050 S research cryostat and kept at an anti-freeze medium for immunohistolochemical (IHC) analysis. IHC was carried out following a standard protocol as previously described (Lu, et al., 2014). Briefly, tissue slices were washed in 1X PBS and blocked in 5% goat serum for 30-60 min. Primary antibodies diluted in 5% goat serum were applied and incubated overnight at 4°C, followed by Alexa Fluor^®^ secondary antibodies (Invitrogen™). The tissue slices were then subjected to extended washing using 1X PBS to reduce background fluorescence before mounting with Diamond Antifade Mountant (Thermo Fisher Scientific, P36962). The following primary antibodies were used: Anti-TH antibody (monoclonal, Sigma-Aldrich, T2928, or polyclonal, EMD Millipore, AB152, both used at 1:500 dilution), rat anti-DAT (EMD Millipore, MAB369, 1:1000 dilution), rabbit anti-synj1 (Novus Biologicals, NBP1-87842, 1:500 dilution), guinea pig anti-synapsin 1/2 (Synaptic System, 106004, 1:500 dilution).

### Cell culture and Transfection

MB cultures (Mani and Ryan, 2009; Pan and Ryan, 2012) and cortical cultures (Mani et al., 2007) were prepared as described previously. Ventral MBs (containing both VTA and SN) or cortices were dissected from P0-1 mouse pups and digested using papain (Worthington, LK003178) or trypsin (Sigma, T1005) supplemented with DNase (Sigma, D5025), respectively. MB neurons were then prepared according to our previously published protocol plated at a cell density of 199,000 cells / cm^2^ and grown in the Neurobasal-A based medium supplemented with GDNF (10 ng/mL, EMD Millipore, GF030). Cortical neurons were plated at 142,000 cells / cm^2^ and grown in the MEM-based medium supplemented with insulin (24 μg / ml, Sigma, I6634) and transferrin (0.1 mg / ml, Calbiochem, 616420). Typically, four P0-P1 mouse brains are required for a MB culture. Calcium phosphate was used for transfection to achieve sparse expression and to ensure analysis of single neurons during the imaging experiments. Transfection was carried out at DIV 3-5 for MB neurons and at DIV 5-6 for cortical neurons, after which, the growth medium was replaced with a fresh medium supplemented with an antimitotic agent, ARA-C (Sigma-Aldrich, C6645).

### Optical setup and live cell imaging

For live cell imaging, cells were mounted on a custom-made laminar-flow stimulation chamber with constant perfusion (at a rate of ~0.2-0.3 mL / min) of a Tyrode’s salt solution containing 119 mM NaCl, 2.5 mM KCl, 2 mM CaCl2, 2 mM MgCl2, 25 mM HEPES, 30 mM Glucose, 10 μM 6-cyano-7- nitroquinoxaline-2,3-dione (CNQX), and 50 μM D, L-2-amino-5-phosphonovaleric acid (AP5) and buffered to pH 7.40. NH_4_Cl solution containing 50 mM NH_4_Cl, 70 mM NaCl, 2.5 mM KCl, 2 mM CaCl2, 2 mM MgCl2, 25 mM HEPES, 30 mM Glucose, 10 μM CNQX, and 50 μM AP5, buffered to pH 7.40 was used to reveal total pHluorin expression for normalizing exocytosis. All chemicals were purchased from Sigma-Aldrich. Temperature was clamped at 30.0 °C at the objective throughout the experiment. Field stimulations were delivered at 10 V / cm by A310 Accupulser and A385 stimulus isolator (World Precision Instruments). A 1 ms pulse was used to evoke single action potentials. Images were acquired using a highly sensitive, back-illuminated EM-CCD camera (iXon+ Model # DU-897E-BV, Andor Corp., CT, USA). Olympus IX73 microscope was modified for laser illumination. A solid-state 488 nm OPSL smart laser at 50 mW (used at 10% and output at ~ 2 mW at the back aperture) was built into a laser combiner system for millisecond on/off switching and camera blanking control (Andor Corp). PHluorin fluorescence excitation and collection were through an Olympus PLAPON 60XO 1.42 NA objective using 525/50m emission filter and 495LP dichroic filters (Chroma, 49002). Images were sampled at 2 Hz with an Andor Imaging Workstation driven by Andor iQ-CORE-FST (ver2.x) iQ3.0 software.

### Confocal Microscopy

An LSM780 upright confocal microscope driven by the Zeiss Zen Black software was used to examine the immunofluorescence in brain slices. Images were acquired at 1024X1024 pixel resolution using single scans by a 10x (for striatal and cortical synj1 expression analysis, Figure 1-Figure supplement 1D-E and Figure 1-Figure supplement 2) or a 63x lens (for nerve terminal analysis, Figure 2). Immunofluorescence of these images was analyzed using ImageJ.

### HPLC lipid analysis and monoamine analysis

Different brain regions were dissected using rodent brain matrices (ASI, RBM-2000C). Flash frozen mouse brain samples were used for lipid extraction, followed by anion-exchange high-pressure liquid chromatography quantification as described previously (Berman et al., 2008; Landman et al., 2006; Zhu et al., 2015 and Cao et al., 2017a). Striatal samples were collected from freshly dissected brains using 2mm reusable biopsy punch (World Precision Instrument, 504529) and flash frozen for further analysis of DA content and two major DA metabolites, HVA and DOPAC, by the Vanderbilt University Neurochemistry Core.

### Behavioral assays

Male *SYNJ1*^+/−^ mice and their littermate controls were tested for general locomotor activity in an open field chamber in a dark room for 60 min. Motor coordination was assessed by accelerated Rota-rod assay. All mice were subjected to 1-hour habituation in the test room with food and water supply prior to testing. Open-field test – each mouse was placed in the center of a 16 x 16-inch chamber equipped with a Versamax monitor system (Accuscan) in a quiet dark room. The mouse horizontal and vertical movements were monitored and recorded for 60 minutes by a grid of 32 infrared beams at ground level and 16 elevated (3 inch) beams. Quinpirole test - mice were divided into two groups for each genotype which were then subjected to peritoneal injection of a D2R agonist quipirole (0.05 mg/kg) or 1x PBS before being placed into the openfield chamber. Movement was recorded for the following 60 minutes in the dark room as described above. Accelerated Rota-rod test – the mouse was placed on a rotating rod with increasing acceleration from 4-40 RPM over 5 minutes. Each mouse was trained for 2 trials before the test. The duration a mouse spent on the accelerated Rota-rod was averaged for consecutive 3 trials spaced by 15 minutes.

### Stereology microscopy

Mice were perfused and fixed as described above (**Cryostat, Immunofluorescence and antibodies** section). MB brain tissues were cryo-sectioned at 40 μm in thickness using Leica CM3050s and stored in antifreeze media containing 30% ethanol glycol, 25% glycerol and 5% PB. For stereological counting, one in every five slices was selected and a total of 8 brain slices were used from each mouse for IHC labeling. Zeiss Axioplan2 was used for tissue slice imaging with a 20X objective, and Stereo Investigator was used for data analysis using the following parameters: frame sizes: 150 μm × 150 μm; grid sizes: 250 μm × 250 μm; top guard zone height: 2 μm and optical dissector height: 8 μm.

### Data analysis and statistics

All statistical tests were performed in OriginPro 8.2, except the Kolmogorov-Smirnov test, which uses a built-in function at http://www.physics.csbsju.edu/stats/KS-test.n.plot_form.html. Descriptive statistical tests were carried out to determine the distribution of the data sets. All data sets conforming to the normal distribution were subjected to two-sample Student’s *t* test or multiple-sample ANOVA test followed by Tukey’s *post-hoc* tests. P values less than 0.05 was considered statistically significant. For human brain data (Figure 1C), the size of each bin was arbitrarily determined to balance sufficient samples and to represent the empirical stages of life (0-10 years: childhood / developmental stage; 11-20 years: adolescent / developmental stage; 21-30 years: early adulthood / full-grown with no degeneration; 30-40 years: adulthood / possible signs of aging and degeneration; 41-60 years: mid-aged / signs of aging and degeneration; 61-70 years: aged / progressive degeneration). For immunohistochemical analysis in Figure 1-Figure supplement 1 D-E and Figure supplement 2, three rectangular regions of interest were placed randomly in the cortical or the striatal region to cover an equal sized area in both regions. Both sides of the hemispheres were analyzed from 2 brain slices for each of the 3 mice at the selected age. For analysis of nerve terminal synj1 immunofluorescence, confocal images were analyzed using the ‘Time series analyzer V3.2’ plug-in by ImageJ. Universal 2×2 μm circular regions of interest were used and placed manually in a double-blinded fashion. Nearly 5% of the measurements that fall outside of the mean ± 3x standard deviation were excluded as outliers. The mean value of immunofluorescence was calculated for each image across 30-80 regions of interest and these mean values were then averaged across all images (10-20) sampled from each animal. Each data point represents the averaged value for one animal and the error bar represents standard error from 10-20 samples. For quantification of DAergic terminals in striatal slices (Figure 5A-B), circular regions of interests were manually placed on all colocalized puncta on the 135 X 135 μm image. To reduce the error of this relatively arbitrary measurement, the following strategies were used: 1) Analysis was performed in a double-blinded fashion. 2) The immunostaining procedure, the confocal imaging settings, and the gain / contrast of the images during analysis were kept the same for a matching number of *SYNJ1*^+/+^ and *SYNJ1*^+/−^ samples. 3) A matching number of images from *SYNJ1*^+/+^ and *SYNJ1*^+/−^ mice were assigned for analysis at a single time. 4) Counts were compared between two analysts. For pHluorin imaging study (Figure 3D-E), data was collected from 2-3 batches of cultures. Each data point represents an average of 3 stable trials on a single cell before and after drug treatment (connected by a black line). Typically, 15-50 nerve terminals with consistent responses were selected for analysis for each cell.

## Acknowledgements

The work was funded by a Cote Early Investigator Award (PD12-00011) to P-Y Pan, an International Research grant (PDF-IRG-1447) from the Parkinson’s Disease Foundation (PDF) to Z Yue and a R21 grant (1R21NS095155-01) from NINDS to P-Y Pan and Z Yue.

---

**Figure 1-Figure Supplement 1.**
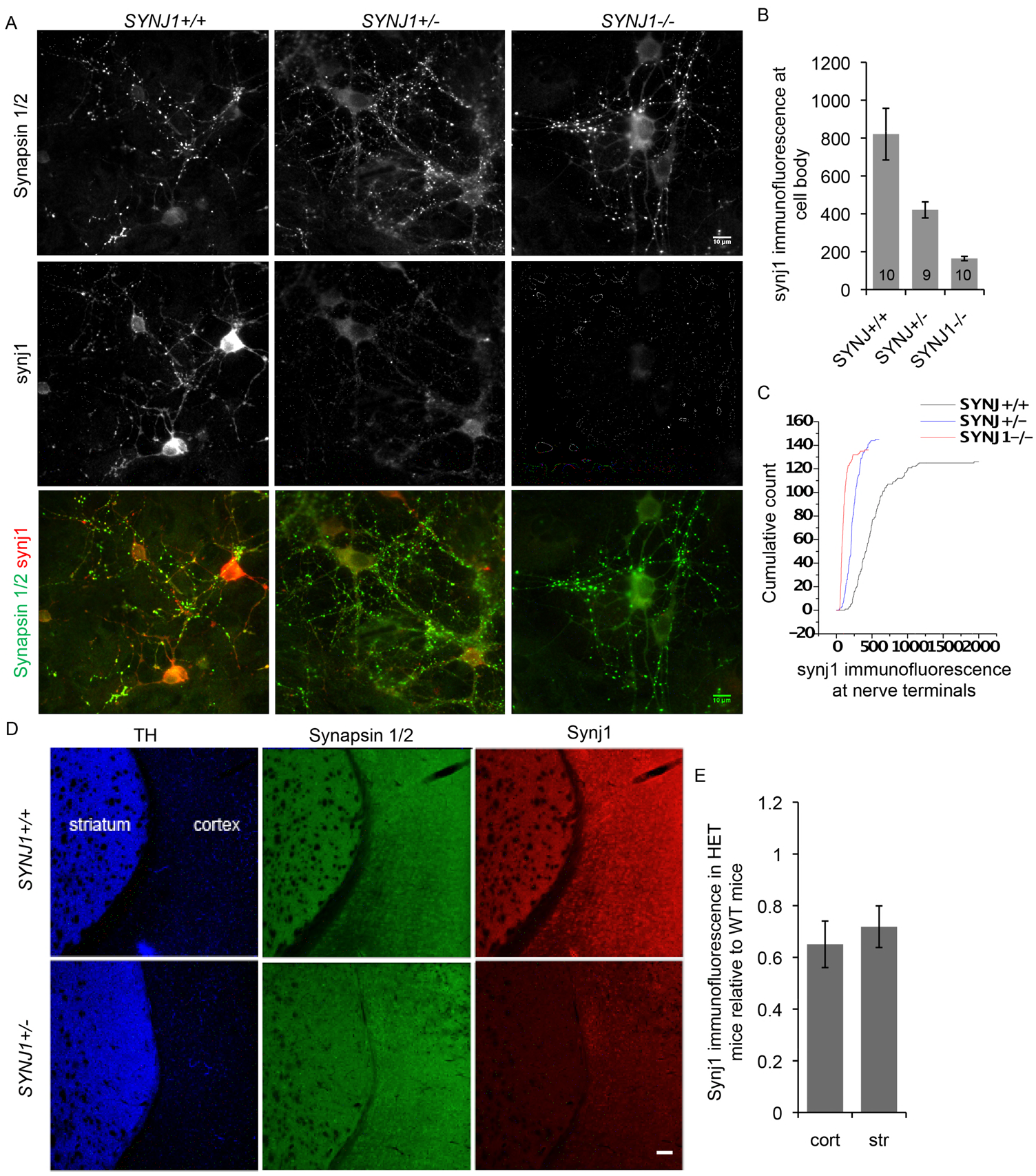
Validation of the synj1 antibody for quantitative analysis. **A**) DIV14 cortical neurons from *SYNJ1*^+/+^, *SYNJ1*^+/−^ and *SYNJ1*^−/−^ mice immunolabeled for synapsin1/2 and synj1. Synj1 display typical nerve terminal enriched expression. Scale bar, 10 μm. **B-C**) Quantification of synj1 immunoreactivity at the cell body (**B**) and at the nerve terminals (C) after background subtraction revealed a gene-does dependent change. **D-E**) Analysis of synj1 immunoreactivity in coronal slices (see Materials and Methods / Data analysis) from *SYNJ1*^+/+^ (N=6) and *SYNJ1*^+/−^ (N=6) mice. Scale bar, 100 μm.

**Figure 1-Figure Supplement 2.**
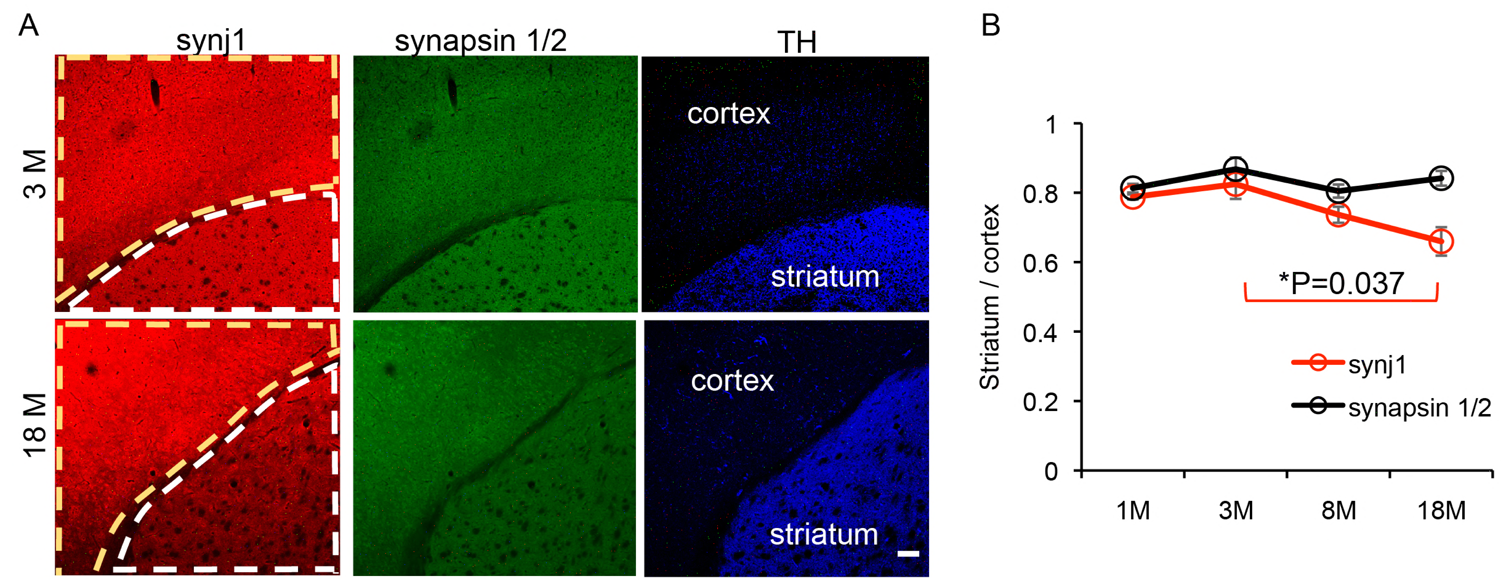
Synj1 immunofluorescence is down regulated in the striatum of aged mice. **A**) Representative immunofluorescence for synj1, synapsin1/2 and TH in the coronal striatal slices of a 3-month old and an 18-month old mouse. Scale bar, 100 μm. **B**) Summary of the expression of synj1 (red) and synapsin1/2 (black) in the striatum (outlined in white in **A**) relative to the cortex (outlined in yellow in **A**) in different ages of the mouse brain (see Materials and Methods / Data analysis). Data represents the average of 3 mice for each age group. To test the null hypothesis that age does not change synj1 immunofluorescence, one-way ANOVA (*P = 0.042 for synj1 and P = 0.29 for synapsin1/2) was performed for each marker. P value represents Tukey’s *post-hoc* analysis following one-way ANOVA.

**Figure 5-Figure Supplement 1.**
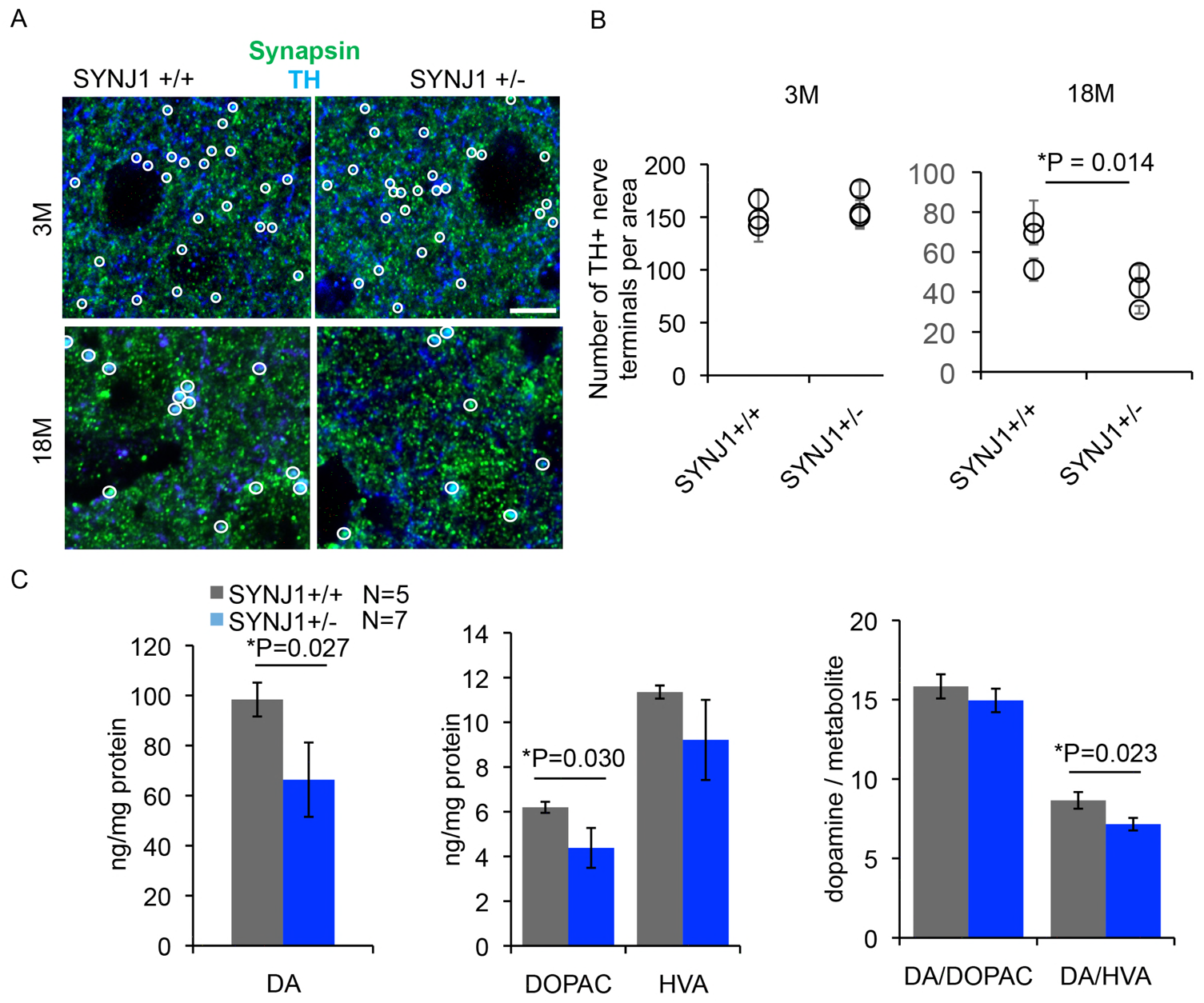
*SYNJ1*^+/−^ mice do not exhibit loss of DAergic cell bodies. A) Representative images showing TH immunolabeling (DAB enhanced, brown) and Nissl counter staining (blue) at the ventral midbrain from a *SYNJ1*^+/−^ mouse and a *SYNJ*^+/+^ littermate. Scale bar, 200 μm. B) Stereological analysis estimating total number of TH and Nissl positive neurons in male *SYNJ1*^+/−^ mice (N = 4) and *SYNJ*^+/+^ littermates (N = 5) as well as the percentage of TH positive neurons did not reveal any difference.

